# Parental provisioning drives brain size in birds

**DOI:** 10.1101/2021.12.19.470191

**Authors:** Michael Griesser, Szymon M. Drobniak, Sereina M. Graber, Carel P. van Schaik

## Abstract

Large brains support numerous cognitive adaptations and therefore may appear to be highly beneficial. Nonetheless, the high energetic costs of brain tissue may have prevented the evolution of large brains in many species. This problem may also have a developmental dimension: juveniles, with their immature and therefore poorly performing brains, would face a major energetic hurdle if they were to pay for the construction of their own brain, especially in larger-brained species. Here we explore the possible role of parental provisioning for the development and evolution of adult brain size in birds. A comparative analysis of 1,176 bird species shows that various measures of parental provisioning (precocial vs altricial state at hatching, relative egg mass, time spent provisioning the young) strongly predict relative brain size across species. The parental provisioning hypothesis also provides an explanation for the well-documented but so far unexplained pattern that altricial birds have larger brains than precocial ones. We therefore conclude that the evolution of parental provisioning allowed species to overcome the seemingly insurmountable energetic constraint on growing large brains, which in turn enabled bird species to increase survival and population stability. Because including adult eco-and socio-cognitive predictors only marginally improved the explanatory value of our models, these findings also suggest that the traditionally assessed cognitive abilities largely support successful parental provisioning. Our results therefore indicate that the cognitive adaptations underlying successful parental provisioning also provide the behavioral flexibility facilitating reproductive success and survival.

**Significance Statement:** The young of large brained species, if left to grow their own brain, would face a seemingly insurmountable energetic constraint, because brain tissue is energetically costly but adequate cognitive benefits arise only after a delay. We therefore hypothesize that protracted parental provisioning was a precondition for the evolution of large brains. Comparative analyses of 1,176 bird species confirmed that parental provisioning strongly predicts variation in relative brain size, suggesting that these two traits coevolved. These results provide the first explanation for the well-known difference in relative brain size between altricial and precocial birds. They also cast doubt on the explanatory value of previously considered social or technological cognitive abilities, suggesting we rethink our approach to cognitive evolution.

## Introduction

Large brains bring diverse cognitive benefits, including enhanced information acquisition and processing (e.g., stereoscopic vision in primates (1, 2) or electro-sensing in mormyroid fishes (3)), motor control and decision making (4-7), and cognitive abilities such as learning (8-10) and reasoning (11-13). This effect is supported by comparative studies examining links between brain size and specific eco-or socio-cognitive abilities (14-19). Comparative studies also indicate that these abilities are adaptive by showing a positive association between relative brain size and longevity in diverse lineages (20-22). On a population level, the cognitive benefits of large brains are reflected in the ability to successfully establish populations (23, 24) and have more stable populations (25).

Nonetheless, despite these cognitive benefits, many species have relatively small brains, which may be explained at least in part by the high energetic costs of brain tissue. The expensive brain hypothesis focuses on the link between relative brain size and the energy available to sustaining the adult brain (26). A number of comparative studies support this idea (27-29). These high energy costs of brains may also have developmental consequences. Developing brains do not yet bring the substantial cognitive benefits required to support the high cost of brain growth and maintenance (30, 31). Thus, an immature of a larger-brained species faces a seemingly insurmountable bootstrapping problem: how can it develop a large, functioning brain when it ideally would already have one to provide the energy necessary for this process? Therefore, when applied to development, the expensive brain hypothesis proposes that natural selection cannot always respond to opportunities to evolve specific cognitive adaptations via an increase in brain size, even if this would in principle lead to higher fitness (32).

This catch-22 leads to the hypothesis that adult brain size across species depends on the parents’ capacity to provision their young (i.e., via egg size, feeding and keeping the young warm (see 32)). It makes similar predictions as the maternal energy hypothesis (33), which argues that the ability of mammalian mothers to transfer energy to their offspring constrains brain size. However, this idea did not consider brain size an adaptive trait, only considered some components of parental provisioning, did not compare its predictive power with that of socio-or eco-cognitive selective agents, and did not recognize the bootstrapping problem (32), and thus failed to derive more detailed predictions. Although various subsequent studies have reported links between separate features of maternal allocation and relative brain size in mammals (33, 34), birds (17), cichlid fish (35), and sharks and rays (36), here we leverage the recently developed parental provisioning hypothesis (32) to provide a comprehensive test that considers all aspects of parental provisioning of young, while at the same time comparing its predictive power for brain size with that of the usually considered eco-and socio-cognitive benefits accruing to adults.

Here we focus on birds, a lineage that is well suited for an integrated test of these different hypotheses. Birds show appreciable variation in many relevant traits, and the energy transfer from parents to offspring can readily be assessed via egg volumes and the time offspring are provisioned. Moreover, birds have a deeply rooted split in the developmental state of the hatchlings, ranging from precocial (no or little provisioning of the young) to altricial (extended provisioning), henceforth referred to as developmental mode (37). Previous work noted that precocial species have smaller relative brain size than altricial ones (38, 39), but the evolutionary causes of this difference remain unclear. Birds also vary in the number of providers (40). These differences allow us to distinguish the effects of investment in eggs from those of provisioning of young. Moreover, birds show major variation in ecology (i.e., aspects of their ecological niche, the climate in their geographic range (41)) and sociality (42).

This high variability in all relevant variables allowed us to simultaneously test their possible effects on adult brain size. We assembled information on adult brain size, adult body mass, predictors assessing parental provisioning (developmental mode, egg mass, clutch size, duration of food provisioning, number of providers), ecology (fiber and energy content of food, complexity of food acquisition, foraging substrate, sedentariness, insularity, climate in the geographic range), and sociality (strength and stability of pair bonds, breeding alone or in colonies, group sizes outside the breeding season) for 1,176 bird species. For the comparative tests, we used phylogenetically controlled Bayesian mixed models (43) to assess the effects on relative brain size across species of i) parental provisioning, ii) eco-social predictors, and iii) all predictors combined. Next, to understand how the different predictors may have influenced each other’s evolution, we used phylogenetic d-separation path analysis (44).

## Results

Because body mass is tightly correlated with brain size, models using absolute brain size leave little variation left to explain. We therefore report the results using relative brain size scaled by body mass as the response variable, but note that analyses including absolute brain size as a response variable produce qualitatively the same results (see Table S1a-c). Moreover, body mass scales with many aspects of bird physiology and life-history in a non-linear manner. Thus, we include body mass and its interaction with key predictors in all models to capture these nonlinearities.

Phylogenetically controlled mixed models showed that larger relative brain sizes in birds were positively associated with all components of parental provisioning (Fig. 1, Tables S2a-d). Larger relative brains were found in species that are altricial, have larger relative egg mass, smaller clutches, and feed their young for a longer time. The effect of these predictors was modified by interactions among them, further highlighting the importance of parental provisioning (Fig. 1). Accordingly, only precocial species showed a negative correlation between relative brain size and clutch size (Fig. 2a), while only altricial species showed a positive correlation between relative brain size and the time offspring are fed (Fig. 2b). The remaining interaction effects reflected fundamental allometries or differences in body mass between precocial and altricial species (Fig. 1). In sum, the total amount of energy invested into each individual young strongly affected relative adult brain size.

**Figure 1.**
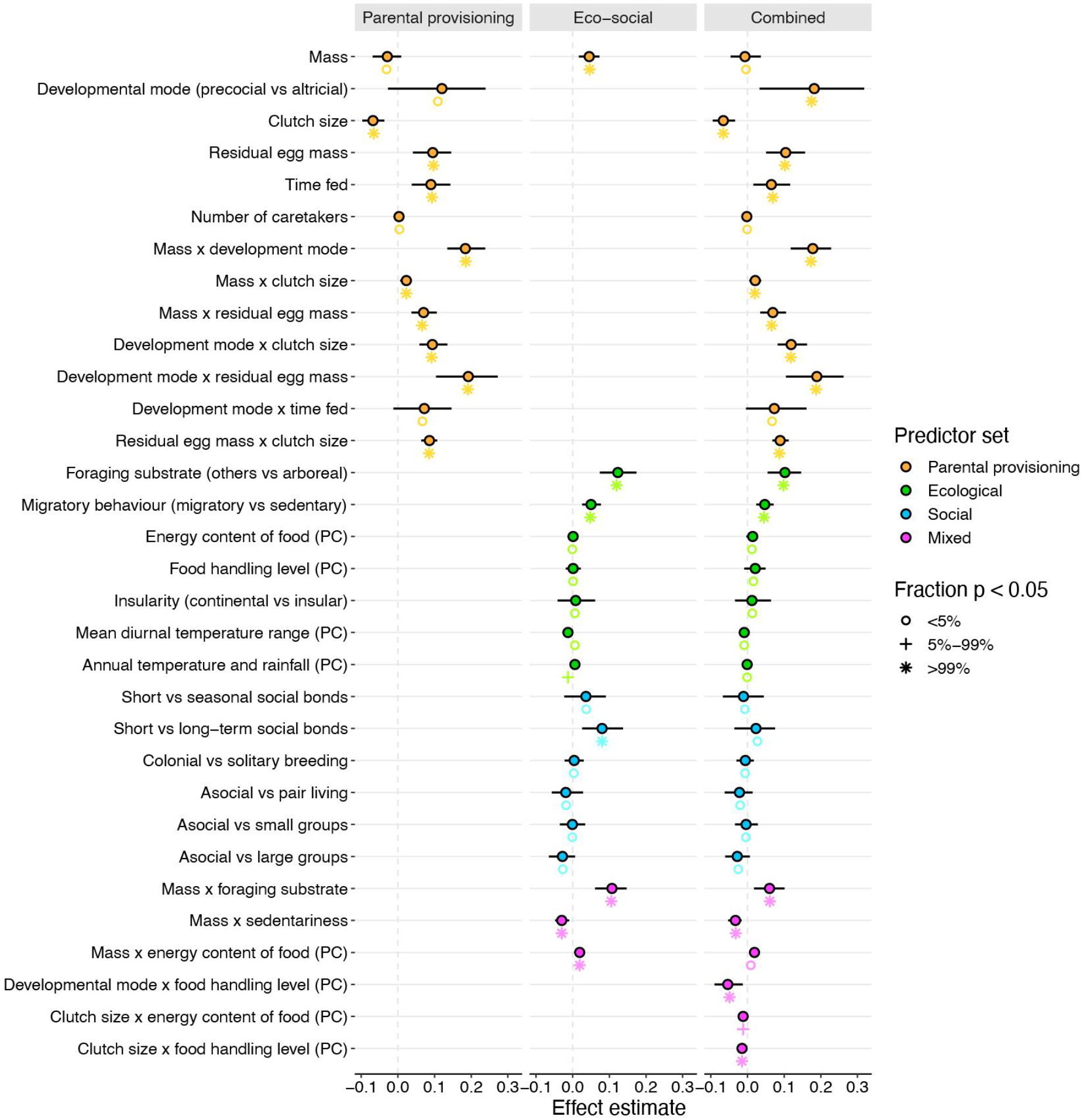
Estimated coefficients and effect sizes of phylogenetically controlled mixed models of the parental provisioning model, the eco-social model, and the combined model on relative brain size in birds. Color-filled circles denote estimated effects, lines denote the 95% lower and upper confidence limits generated in the R package MCMCglmm (43) based on a consensus phylogeny. Symbols below the circles denote the proportion of random tree models that reach statistical significance (p<0.05) for each predictor based on model averaging of 200 models, using a set of random trees from http://birdtree.org (78). The corresponding full models are shown in S2a-c Table.

**Figure 2.**
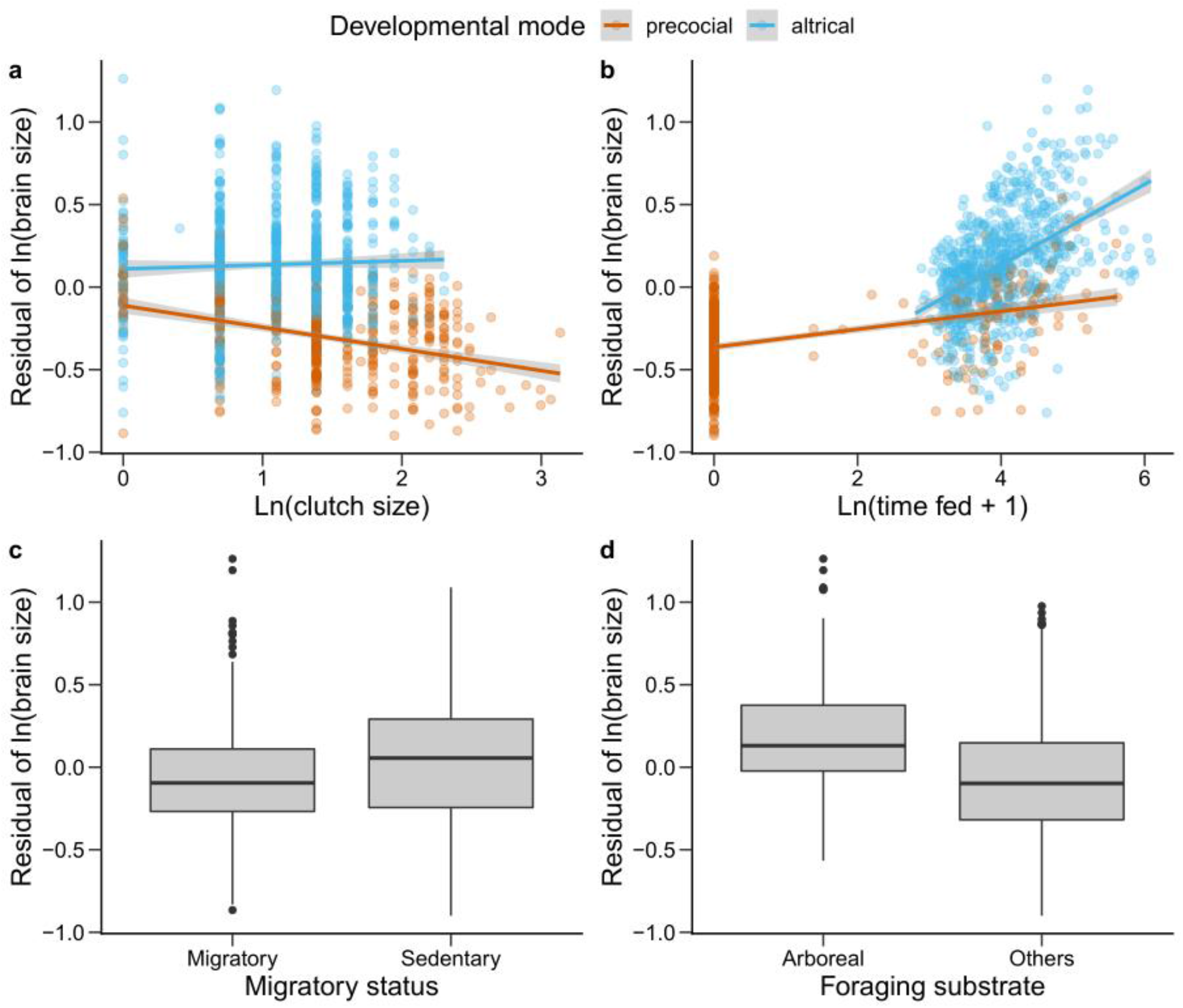
The effect of parental provisioning (a, b) and ecological predictors (c, d) on relative brain size in birds (N=1,176 species). **a)** Clutch size vs developmental mode (precocial (N=427 species) vs altricial (N=749 species)). **b)** Time provisioned vs developmental mode (precocial vs altricial). **c)** Movement pattern: sedentary (N=566 species) vs migratory (N=610 species). **d)** Foraging substrate: others (N=929 species; including terrestrial, aerial, aquatic foraging) vs arboreal (N=247 species). In **a-b**, solid lines designate linear trend estimates, grey shading designate standard error bands. In **c-d**, thick lines designate medians, the grey boxes designate inter-quantile ranges (IQR, 0.25-0.75 quantile), whiskers designate ±1.5 IQR, circles designate outliers.

In a separate analysis of the effect of eco-social predictors on relative brain size, phylogenetically controlled mixed models primarily showed that sedentary species and those that forage arboreally, had larger relative brain size than those that are migratory or use other foraging substrates (Fig. 2c,d). In terms of interaction effects, we found that relative brain size increased with body mass only in arboreal foragers, and that higher energy content of food translated into larger relative brain size only in larger species (Fig. 1). Social factors had a much weaker impact on relative brain size: species with enduring pair bonds tended to have a larger relative brain size than species without pair bonds, whereas no correlations were found for the other social predictors (Fig. 1).

This eco-social model explained markedly less variance in relative brain sizes than the parental provisioning model (R^2^ = 0.38 vs R^2^ = 0.50). However, a combined model containing both parental provisioning and eco-social predictors performed best (R^2^ = 0.57). Although confirming the patterns of the separate models (Fig. 1), it revealed that the posterior estimates of effect sizes of parental provisioning predictors were far larger than those of the eco-social factors. Among the latter, the ecological predictors that remain significant in the combined model were largely linked to the adult foraging niche and movement patterns, whereas social predictors no longer played any role.

Additional models showed that data limitations are not responsible for the observed patterns. Models that excluded predictors with limited data availability (time fed, social grouping; Table S3 a-b) confirmed the key role of parental provisioning in relative brain size. Similarly, using a more detailed categorization of the developmental mode spectrum instead of binary categorization recovered the same patterns (Table S3c), and so did a model that controlled for the possible confounding effect of longevity on parental provisioning predictors (Table S2d). Finally, lineages may differ in their brain-to-body size allometry (45), which may affect the patterns. However, a model using lineage-specific (46) brain-to-body allometries (Table S4) confirmed the results of the analyses using the overall allometry.

The effect of parental provisioning on relative brain size became obvious when superimposed on a phylogenetic tree (Fig. 3). Starting at 3 o’clock and moving counter-clockwise, early emerging, precocial lineages had relatively small brains (Palaeognathae ❶, Galloanserae ❷, Charadriiformes ❹), while inter-dispersed altricial lineages (Columbimorphes ❸, Suliformes, Pelicaniformes, Procellariiformes ❹, Strisores ❽) had slightly larger brains. With the origin of Inopinaves (core land birds: Accipitiformes-Passeriformes ❾-⓮) some 65 million years ago (46), systematic altriciality and extensive parental provisioning arose, resulting in several clusters of lineages with notably enlarged relative brain size, including owls, parrots, and corvids, while one lineage (Strisores ❽) reverted to precociality and small brains. The most recently evolved and highly diverse other passerines were small-bodied and did not have brains of exceptionally large relative size, although they are still relatively larger than those of precocial lineages.

**Figure 3.**
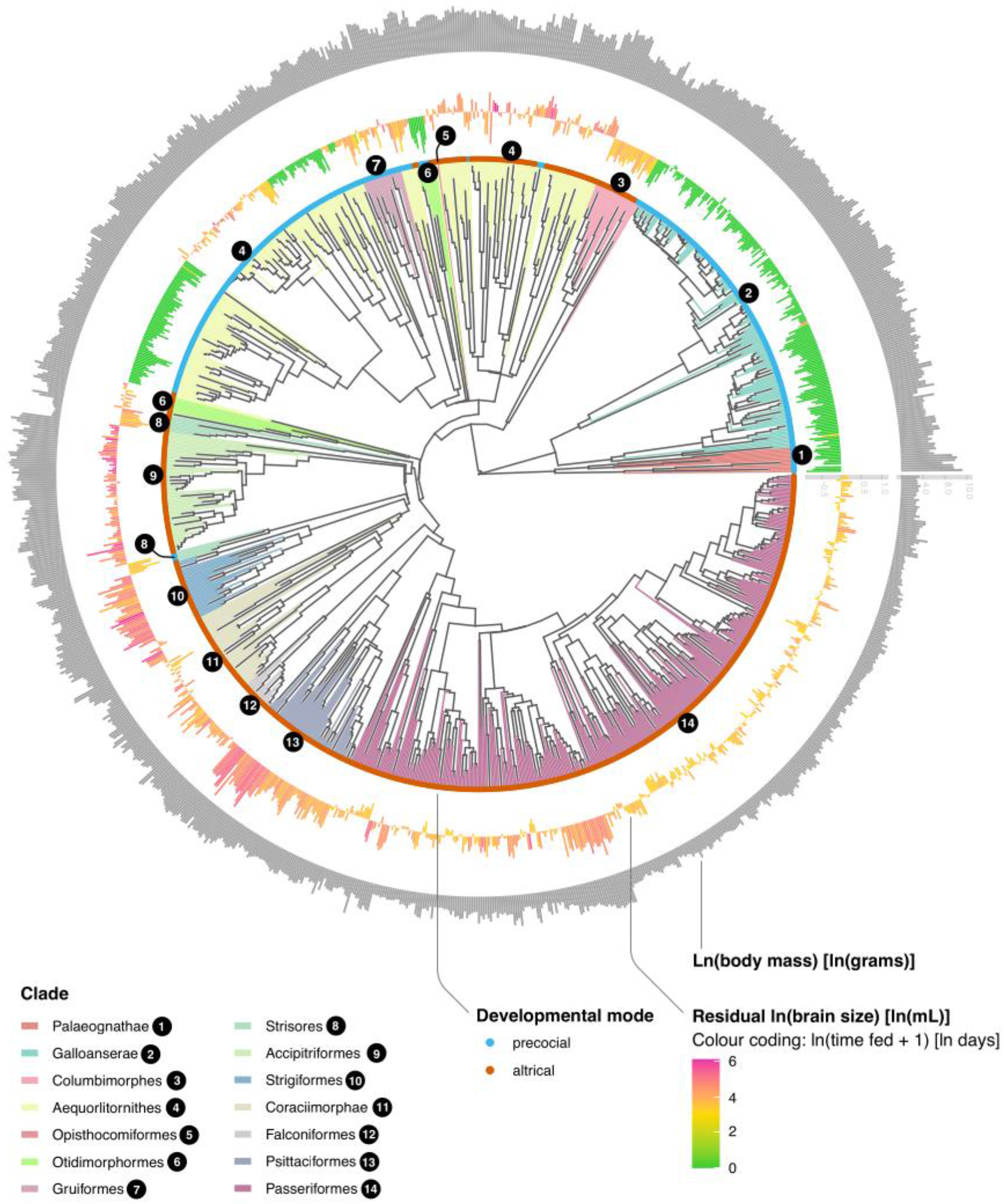
Phylogenetic distribution of relative brain size, developmental mode (precocial vs altricial), and time parent(s) feed their offspring (N=1,176 bird species). Size of bars in outer circle represent body mass (ln(grams)); size of bars in middle circle represents relative brain size (ln(mL)) while their color represents the time parents feed their offspring (ln(days + 1)); inner circle represents developmental mode (precocial, altricial). Phylogeny follows birdrtree.org (78), whereas taxonomy largely follows Prum et al. (46), explaining why some clades appear polyphyletic.

Because the predictors analyzed above may not only have affected relative brain size directly, but may also have selected for evolutionary changes in values of other predictors and thus affected brain size indirectly, we performed phylogenetic path analyses (44, 47) to gain better insights into the evolutionary relationships between traits. We included well-established developmental and eco-social relationships in all models, but also added biologically plausible ones involving brain size that were varied across the different models.

The model assessing the relationships among the various parental provisioning predictors of relative brain size showed that adult body mass affects all other traits, including developmental mode (Fig. 4a). Both a large adult body mass and altriciality had a negative effect on clutch size and residual egg mass. Altriciality had a strong positive effect on the time offspring are fed, and subsequently on brain size. These patterns reflect established life-history trade-offs and their cascading effects on the total amount of energy transferred from parents to their offspring at the different developmental stages. The remaining effect of body mass on relative brain size depended on developmental mode, reflecting that larger altricial species had larger relative brain sizes than precocial species, relative to what would be predicted from body size alone (see Fig. 1). This pattern therefore fully supports the pervasive effect of parental provisioning on a species’ brain size.

**Figure 4.**
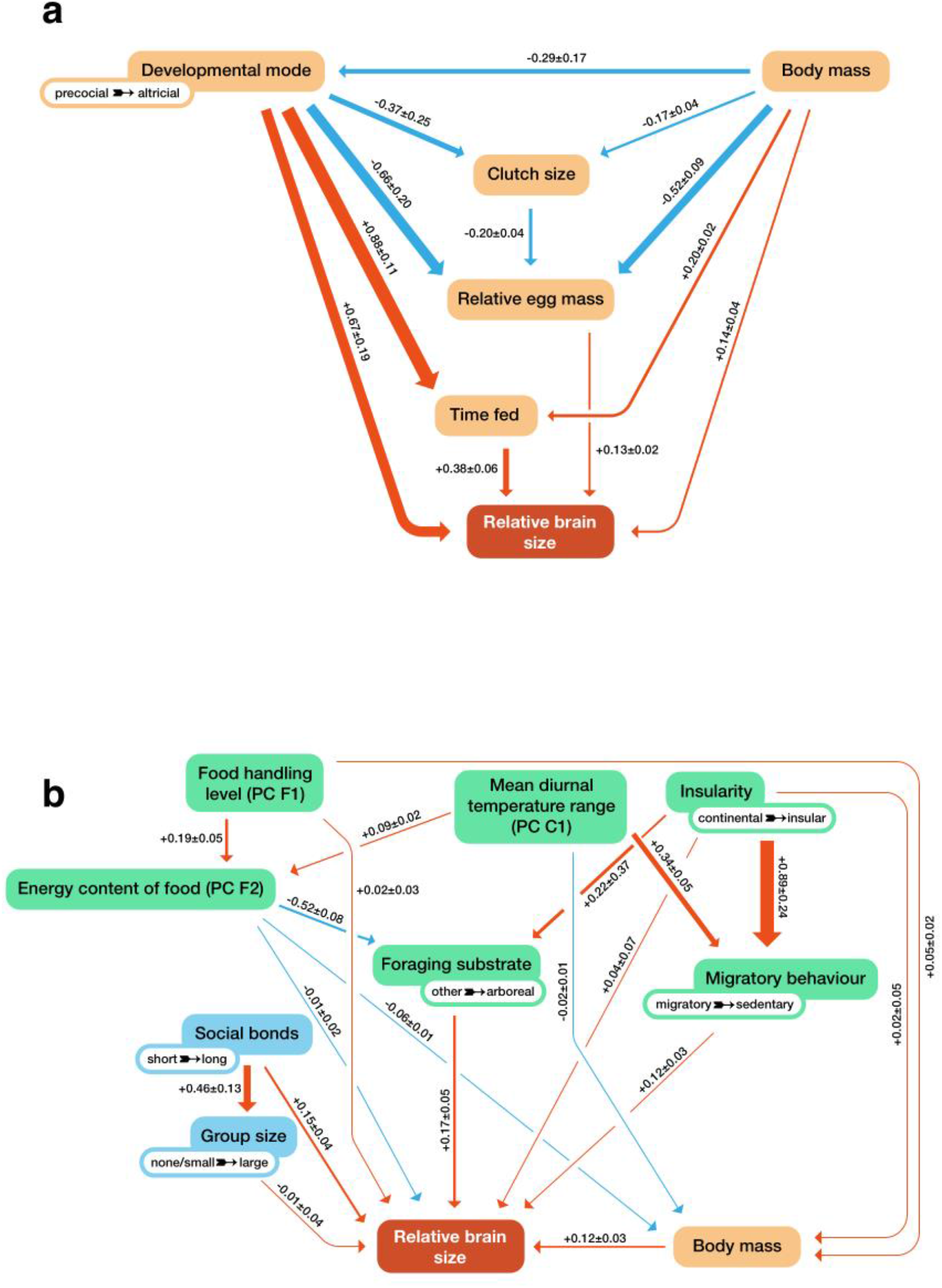
Phylogenetic associations among specific predictor sets and relative brain size, assessed by d-separation path analyses (44) (N=1,176 bird species). Arrows thickness and numeric values show the strength of the association, arrow color shows its direction (positive: orange, negative: blue). **a**, Associations among parental provisioning predictors. **b**, Associations among eco-social predictors.

Turning to the eco-social predictors, the path model assessing the evolutionary relationships among them showed that some of these predictors have strong evolutionary links among themselves, but rather weak selective effects on relative brain size (Fig. 4b). We note that a model where social predictors and the foraging substrate were a consequence of larger brains did perform significantly less well than the model where these predictors were linked to relative brain size. A combined path analysis (Fig. 5) including both parental provisioning and eco-social predictors retained the relationships revealed by the separate and combined phylogenetically controlled mixed models. Parental provisioning continued to have a direct and strong effect on relative brain size, whereas that of the eco-social variables remained weak.

**Figure 5.**
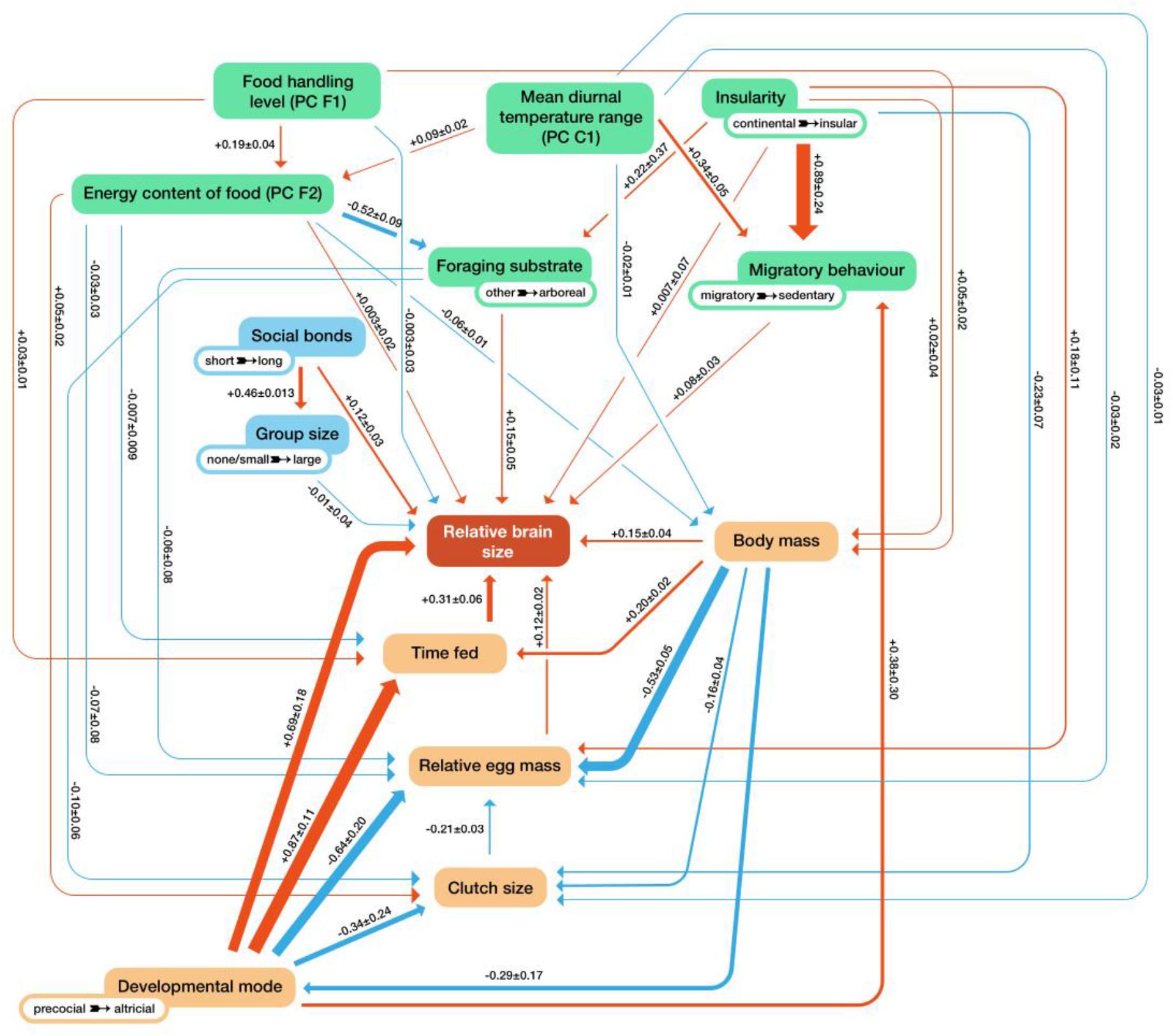
Phylogenetic associations among parental provisioning (orange), ecological and social predictors (green) and relative brain size, assessed by d-separation path analyses (44) (N=1,176 bird species). Arrow thickness and numeric values show the strength of the association, arrow color its direction (orange: positive, blue: negative).

## Discussion

For animals, developing and maintaining brains, particularly large ones, is energetically demanding (30, 31) during a time when they are still developing the requisite cognitive abilities to acquire the energy. This creates a bootstrapping problem. The parental provisioning hypothesis (32) highlights the pivotal role of the energy invested by parents into the development of their young in overcoming this bootstrapping problem. Large brains can therefore only evolve in species where parents are capable of sustaining both themselves and their growing young, and this process is supported by the cognitive abilities large brains produce. Consequently, the proximate and ultimate drivers of brain size are tightly linked and no clear distinction between them is possible (32).

The analyses reported here confirm the predictions of the parental provisioning hypothesis, and the results have three major implications, which we now discuss in turn. First, parental provisioning is the main predictor of adult relative brain size in birds: in the combined model, adult eco-and socio-cognitive demands hardly play a role. Thus, large brains can only be sustained in species with extensive energy inputs during development. This association is supported by previous work in mammals (34), birds (17), fish (35), as well as sharks and rays (36). However, these studies overlooked the bootstrapping problem faced by developing young (33, 34). The parental provisioning hypothesis offers a novel theoretical underpinning to this pattern (32). Parental provisioning enables growing young to construct a large brain during a life stage when they themselves have not yet learned the skills needed to acquire the energy necessary to sustain such a growing and learning brain. Accordingly, parental provisioning removed this bottleneck for the evolution of larger brain size, and enabled bird species to evolve into skill-intensive niches, as evident by the diverse foraging and nesting habits of altricial species compared to precocial species (48). The importance of parental provisioning is also highlighted by the phylogenetic path analyses, which confirm that its components form a tightly coevolved set.

Second, the parental provisioning hypothesis provides the first explanation for the well-established pattern that altricial species have relatively larger brains than precocial species (38, 39), and that altricial species have steeper brain-body allometric slopes (32, 49). Altricial young are provisioned by their parents until they have reached adult size. In contrast, most precocial young are only provisioned through egg mass, which is subject to obvious physical limitations. As a result, precocial species experience serious constraints on the evolution of relative brain size and therefore niche complexity (48). The evolutionary transitions to altriciality allowed species to evolve into cognitively more demanding niches, which require more elaborate food-handling or anti-predator skills (39). This was made possible by parents donating the requisite large brain to their offspring through extended parental provisioning. However, these transitions were actually quite rare (39). Although the underlying reasons for this rarity remain unclear, we speculate that precocial species remain stuck in a simple ecological niche (48) unless they find a way to breed in safer places, which facilitates the evolution of provisioning and therefore altriciality.

Finally, our findings imply a modest role at best for socio-and eco-cognitive abilities in the evolution of brain size in birds. We found almost no detectable effect of the cognitive demands of adult social life. We did, however, find weak evidence for effects of eco-cognitive predictors independent of parental provisioning that affect the energy balance of adults. First, migratory species have smaller brains than resident species (50), indicating that the high energy cost of migration reduces the ability to sustain a large adult brain. Second, the effect of arboreal foraging may be more related to adult survival, as in mammals (51), thus removing a constraint on brain enlargement.

The absence of an effect of the traditionally measured socio– and eco-cognitive variables on relative brain size is puzzling. It may actually reflect a more general problem in comparative cognition, as popular tests often fail to produce a differentiation between species generally thought to show major differences (52). Thus, these tests may not assess those cognitive adaptations that produce fitness benefits by scaffolding critical foraging and predation-avoidance skills and improving the ability to provision the young and/or improve survival. Our results suggest we should pay closer attention to cognitive adaptations such as general behavioral flexibility (as enabled by executive functions and domain-general intelligence (13)), including the coordination of parental duties (essentially linked to planning, decision-making, and time management). These cognitive abilities may be expressed behaviorally in avoidance of predation on eggs, nestlings and adults (53, 54), in maximization of food intake under everchanging food and weather conditions, or in the socio-cognitive ability to coordinate offspring provisioning (14). For example, birds can flexibly adjust parenting in response to predation risk to themselves and their nestlings (55), as confirmed experimentally (56). However, because these cognitive abilities are not readily estimated in a way that can be compared across many species, they are so far not included in comparative analyses of brain size evolution.

To conclude, the amount and duration of parental provisioning is the key to understand brain size evolution in birds, and likely also more generally (32). Across animal lineages, the evolution of parental provisioning beyond eggs (i.e., altricial birds, mammals, cartilaginous fishes) co-evolved with major shifts in brain size (32). Moreover, the increase in the number of caregivers during human evolution and the provisioning children well into adolescence (57) may have largely enabled the massive increase in relative brain size in our lineage over the last 2.5 million years (58).

## Materials and Methods

### Data collection

We gathered information quantifying parental provisioning and the socio-ecological correlates identified by previous work as influencing brain size in animals. We collected these data from the online version of the Handbooks of the Birds of the World (59), region specific handbooks (60, 61), existing published datasets (42, 62-64), as well as species specific studies listed in the dataset. Data on brain size were collected from three sources (19, 65, 66).

The following parental provisioning predictors were included: adult body mass, developmental mode, egg mass, clutch size, the time offspring are fed by their parents until nutritional independence, and the number of caretakers. We did not use imputed data based on mean values of related species, which limited our sample size compared to other comparative studies. Adult body mass data were calculated as means if mass of males and females were listed separately. The developmental mode was assessed dichotomously. Precocial species are those which hatch with open eyes and abandon the nest within 2 days after hatching, and here also encompass semi-precocial species (young hatch with open eyes and are soon capable of walking, but they remain in the nest to be fed by parents). Altricial species are those in which young hatch with closed eyes and largely lack feathers and are fed by parents in the nest, and here also encompass semi-altricial species (young hatch with open eyes and are feathered, but remain in the nest to be fed by their parents) (39). Given that this dichotomous split could obscure patterns, we did re-run the main analyses using all four categories (fully precocial, semi-precocial, semi-altricial, fully altricial; see Table S3c). Egg mass was either assessed through direct measurements of egg mass, or was calculated based on egg dimension measurements, using the formula by Hoyt (67). For those species where multiple egg dimension measurements were available in the literature, the mean values of all provided measurements were used. For those species where egg dimension or mass data were not available in handbooks, we used information in published data compilations listed in the dataset (64, 68-70). Clutch size data corresponded to the median clutch size (54, 70). The time young are fed by their parents included both the time parents provision their young in the nest as well as thereafter until young are nutritionally independent from their parents, as reported in handbooks (see above), and published datasets (38). We also included the mean number of caretakers, where mound-nesting species had zero caretakers, species with uniparental care one, species with bi-parental care two, and cooperatively breeding species the mean value of caretakers. These data were obtained from handbooks (59), existing datasets (40, 66), and species-specific studies listed in the dataset.

Based on previous studies on the drivers of brain size in vertebrates, the following ecological predictors were included: food energy, fiber content of food, number of processing steps required to extract food, foraging substrate, sedentariness, insularity, and climatic predictors. The predictors relating to food were calculated based on a published database (62). We calculated the mean energy and fiber content of foods per 100g from previous studies (71, 72), as well as the number of processing steps required to extract food (73). Given that these three predictors are highly correlated (i.e., low caloric food usually has a high fiber content and requires only a single processing step; Fig. S1), we used a PCA approach and extracted two independent PCs from these predictors (see below). The foraging substrate data were obtained from a published database (63). We categorized species into those that forage arboreally, compared to all other foraging substrates. In mammals, arboreal foraging has been shown to be associated with increased longevity, which in turn allows the evolution of larger brains (51). Sedentariness was assessed based on the maximum movement pattern of a species, separating sedentary species (including local movements) from migratory species (short-and long-distance migrants, altitudinal migrants), combining data from three sources (42, 63, 65). In addition, we also included data on the insularity of species, distinguishing continental from insular species (18, 63). Missing data were directly obtained from the primary sources listed above. Finally, we included data on the climate in the species’ geographic range (74, 75), using a PCA approach (see below). Values of species with contradicting values in different published datasets were confirmed based on the literature listed above, using the above definitions.

The social predictors were also based on previous studies and included social bond strength, group size, and solitary vs colonial breeding. Social bond data were based on published data (76), separating species with short bonds during mating only, seasonal bonds, and long-term bonds that extend beyond a single breeding season. The grouping patterns of species outside the breeding season distinguished between asocial species (usually on their own), pair living species (usually in pairs), species that live in small groups (usually 3-30 individuals), and large groups (usually more than 30 individuals). The split between small and large groups should capture the difference between personalized groups where group members know each other individually, including family groups (42), and ephemeral and/or anonymous groups. We also included whether species breed singly or whether they breed in colonies.

### Analyses

All statistical analyses were carried out in the R 4.0.2 environment (77). The analyses were structured as a series of linear models with increasing complexity, all including phylogenetic relatedness across species (in the form of a phylogenetic correlation matrix based on well-resolved avian phylogenetic trees (78)). For predictors that are expected to be allometrically linked to body mass (brain size, egg mass), the models included their residuals from regressions against body mass. Brain size (the response variable) and adult body mass were log-transformed prior to extracting relative brain sizes. All continuous predictors were mean-centered and scaled to a standard deviation of one, to aid in the interpretation of intercepts and the comparison of effect sizes. The three predictors relating to food (food energy, fiber content, food handling levels) were strongly collinear (Fig. S1a-d, Table S5a). Thus, we used a Principal Component analysis in the package psych (79) (using varimax rotation) to reduce these predictors to two uncorrelated components. Based on their loadings, the resulting two components describe the energy content of food and the food handling levels (including the absence of fibers) (Table S5a). Using the same approach, we extracted three uncorrelated components from 14 climatic predictors. PC1 reflects a latitudinal gradient from the tropics to high latitudes. PC2 reflects inter-annual variability. PC3 reflects a gradient from dry, open and highly seasonal environments to stable and closed environments (Table S5b).

We fitted three phylogenetically controlled mixed models that assessed the dependence of relative brain size on provisioning predictors (model 1), eco-social predictors (model 2), and a combination of both (model 3). Each model included a set of main effects and first-order interactions with meaningful biological interpretation (see Tables S1-4 for a full list of predictors and interactions in each of the models). Since the size of models (both in terms of data and numbers of predictors) prevented a full search of all predictor combinations via selection based on information criteria (IC), we conducted backward simplification of models, removing non-significant interactions (starting with the ones having the largest prediction error). To remain conservative, we did not remove non-significant main effects. Backward selection has been shown to perform well in comparison to IC-based model selection in similar contexts (80), and is robust in cases where the number of individual observations exceeds several fold the number of estimated predictors, as is the case here. We report final outputs of the most parsimonious models (Fig. 1, Tables S1-4). All models were fitted using an MCMC-based mixed modelling approach using the package MCMCglmm (43). We used chains of 50’000 iterations, with the first 10’000 iterations discarded as burn-ins and thinned every 40 iterations. Visual inspection of the final MCMC samples did not show any sign of autocorrelation. All effective sample sizes were close to the actual numbers of sampled draws from predictors’ posteriors (see Tables S1-4). In all models, inverse-gamma priors were used for residual variances (parametrized as inverse-Wishart with V = 1 and ν = 0.002). The prior for phylogenetic effect was formed as a weakly informative half-Cauchy density (parameter expanded priors with V = 1, ν = 1, αμ = 0 and αV = 10’000). Priors for fixed effects were left as default (Gaussian densities with μ = 0 and large variance). Using a modification of the variance inflation factor (VIF) analysis adjusted to the phylogenetic comparative mixed model used, we verified that none of the other predictors were significantly collinear, as all VIF values were <2 (Table S6).

The necessity of complete information on all predictors reduced the sample size of the main models to 1,176 species. Thus, we fitted additional models excluding those predictors that most strongly limited our sample size: excluding duration of parental provisioning (N=1,458 species; Table S3a); excluding duration of parental provisioning and social grouping (N=1,594 species; Table S3b). Both models recaptured the patterns observed in the full model (Table S2), confirming that our sample was not biased by limited data.

The brain-to-body size allometries have been shown to differ across lineages, partly reflecting differences in overall body size across them (45) and partly reflecting the lower parental provisioning in precocial ones (39). These differences may affect our analyses as the residual brain size of larger species in lineages with overall more shallow brain-to-body size slopes will be increasingly overestimated when using an overall Aves-wide allometry to estimate it. Thus, we did re-run the key models using lineage-specific brain-to-body allometries to estimate their residual brain sizes. We used the following monophyletic lineages: Accipitriformes, Aequorlitornithes, Columbimorphes, Coraciimorphae, Falconiformes, Galloanserae, Gruiformes, Opisthocomiformes, Otidimorphormes, Palaeognathae, Passeriformes, Psittaciformes, Strigiformes, Strisores (see 45).

Subsequent to the phylogenetically controlled mixed models, we conducted phylogenetic path analyses to assess the evolutionary links among the predictors included in parental provisioning, the eco-social and the combined model. For each model we defined a set of d-separation statements (44, 47, 81), describing a hypothesis represented by a directed acyclic graph, linking all involved predictors with an assumed causal links. Given the large number of predictors used in the phylogenetic mixed models, we built the different models using three criteria: i) we grouped the variables into social, eco-climatic and life-history traits; ii) we fixed certain relationships between variables based on established patterns (e.g., the island rule linking body size with island/mainland life-style, body mass link with the developmental mode); and iii) we explicitly varied only those paths that directly reflected our hypotheses. Thus, the sequence of models considered reflects hypotheses related to brain size predictors, as outlined in Fig. S2-4. We considered mostly models where brain size is a consequence of eco-social predictors, but our set included also models where social predictors and the foraging substrate/food energy content where a consequence of brain size.

The models were implemented as sets of phylogenetic linear mixed models fitted in the ASReml-R package (82). In models involving binary responses (e.g., precocial/altricial), we used generalized linear mixed models with probit link function to fit models, which makes their regression coefficients directly comparable with standardized regression coefficients from general linear models for continuous predictors. Since all continuous predictors were scaled to unit standard deviation, results for categorical predictors can be interpreted as standardized Cohen’s d effect sizes, and thus, can be directly compared with continuous standardized regression coefficients. The best-fitting hypothesis in each model set was determined via comparisons of their CICc (corrected Fisher’s C Information Criterion, a path analysis analogue of AICc (47)). In the parental provisioning and eco-social model, the best model was clearly supported over the alternatives (ΔCICc > 2 for top models). In the combined model, two models were clearly supported over the alternatives, but were conceptually identical (Table S7).

## Acknowledgments

We thank all field biologists who collected all the natural history data underlying this study. We thank Uri Roll for sharing climatic data, and three anonymous reviewers as well as Jeroen Smaers and F. Stephen Dobson for insightful comments on earlier drafts. This work was supported by a Heisenberg Grant nr. GR 4650/2-1 by the German Research Foundation DFG to M.G., the Polish National Science Centre, grant no. UMO-2015/18/E/NZ8/00505 to S.M.D., the Australian Research Council DECRA award no. DE180100202 to S.M.D., the University of Zurich to C.P.S, and a Max Planck Fellowship to C.P.S.

## Supporting Information

**Fig. S1.**
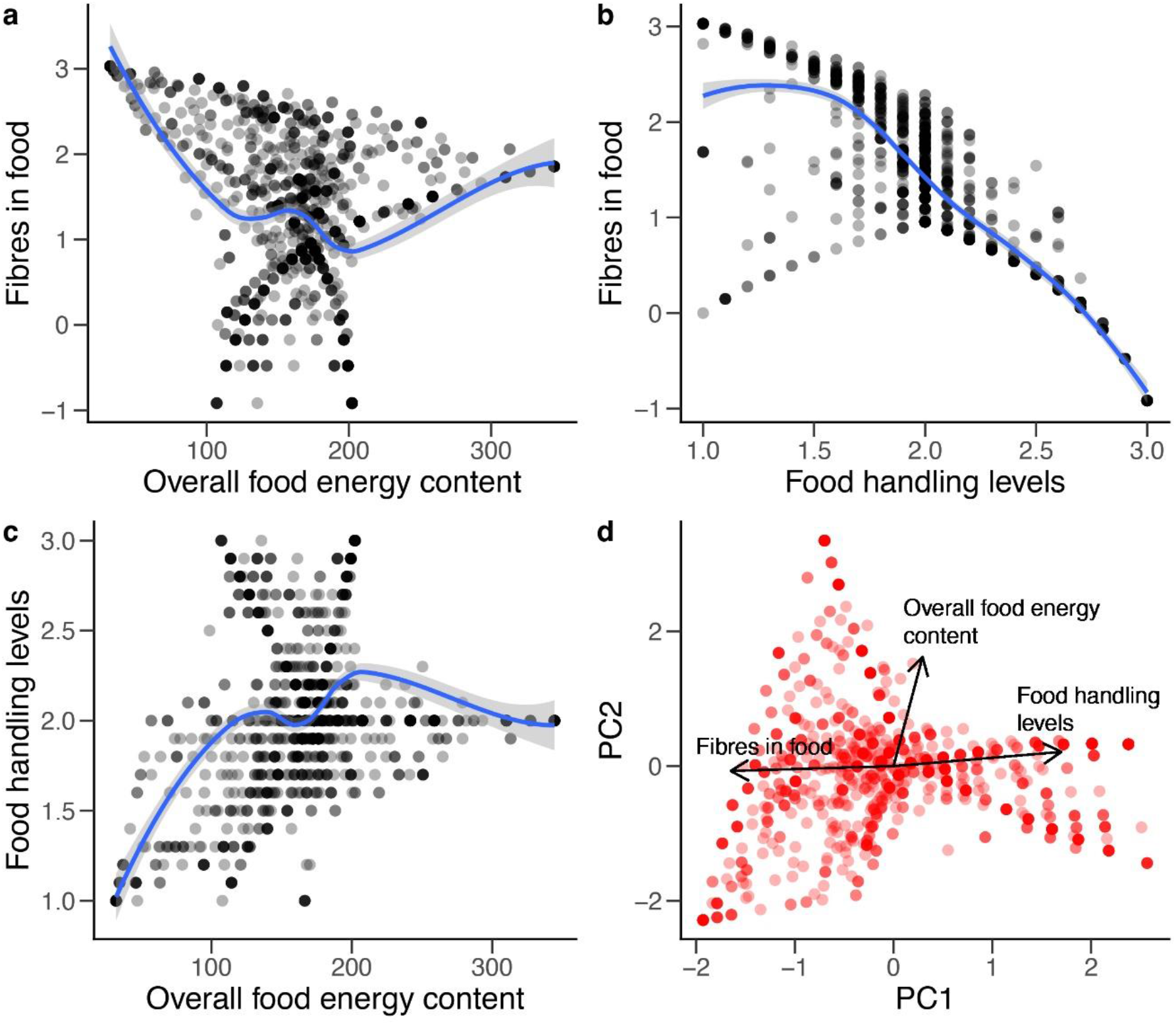
Principal Component Analyses of food related parameters and the variance explained by each factor. **a-c**: bivariate scatter plots showing relationships between the three food parameters. **d**: PCA biplot of the first two PC components. In **a-c**, raw data were augmented by adding a LOESS smoother to approximate the trend (blue lines with shaded standard error bands). In **d** black arrows represent loadings of original variables on the first two PCs. Points designate extracted principal component values (rotated coordinates of each observation for the first two PCs). In **a-d** overlapping points were made semi-transparent to better enable identification.

**Table S1a-c.**
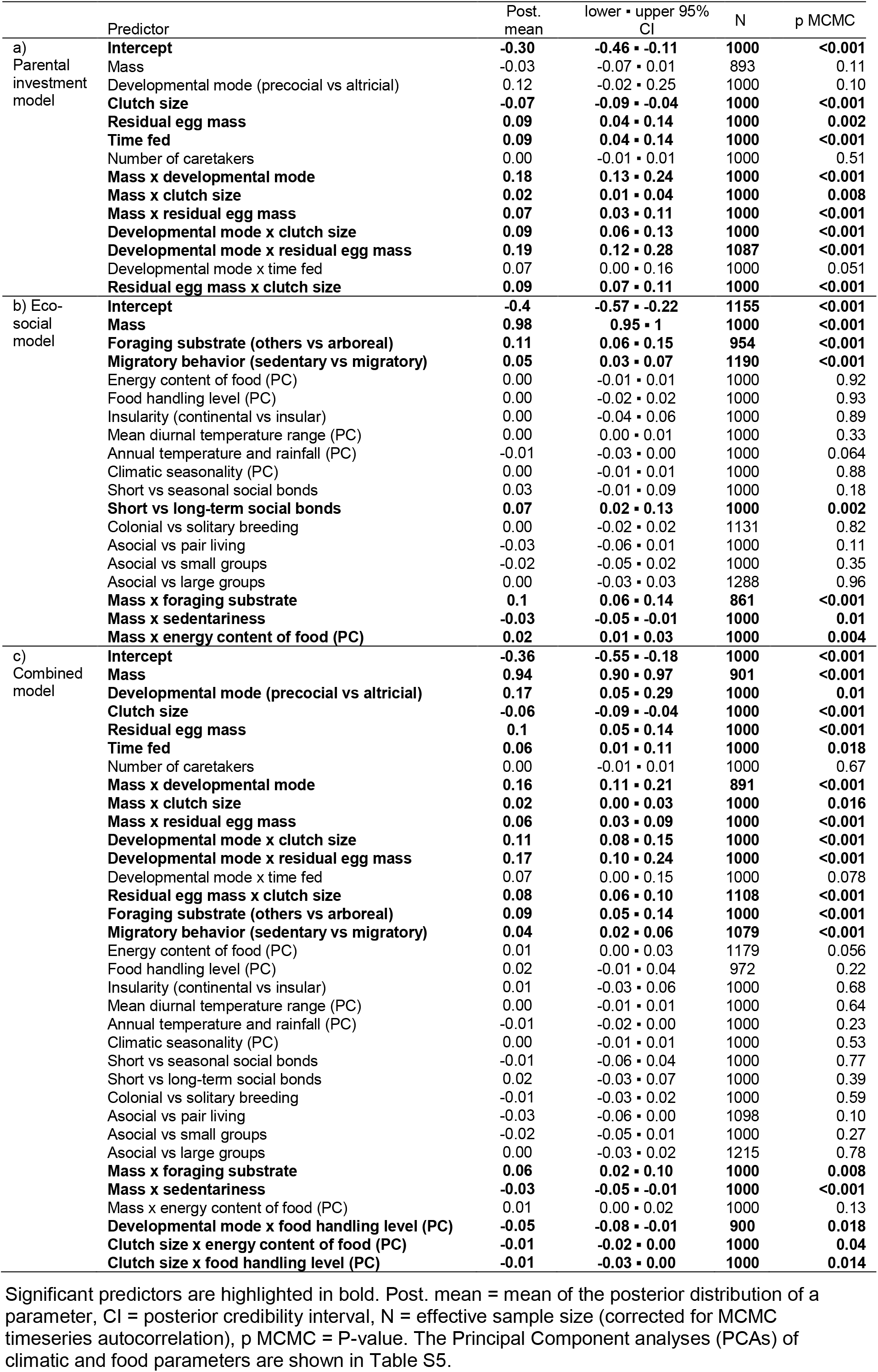
Phylogenetically controlled mixed models in MCMCglmm (1) assessing the effect on absolute brain size in birds (N=1,176 species) of a) parental provisioning predictors, b) eco-social predictors, and c) both combined.

**Table S2a-d.**
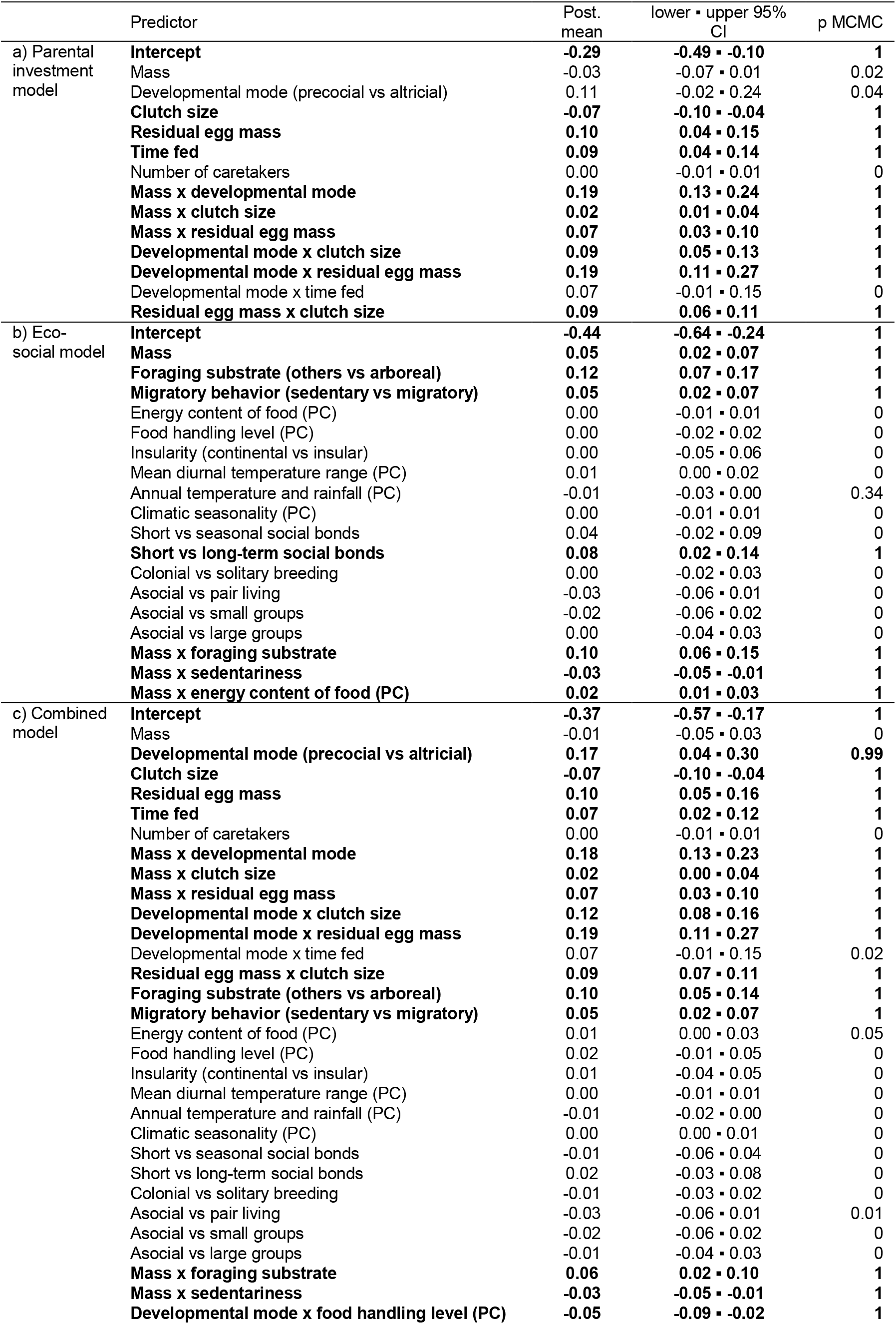
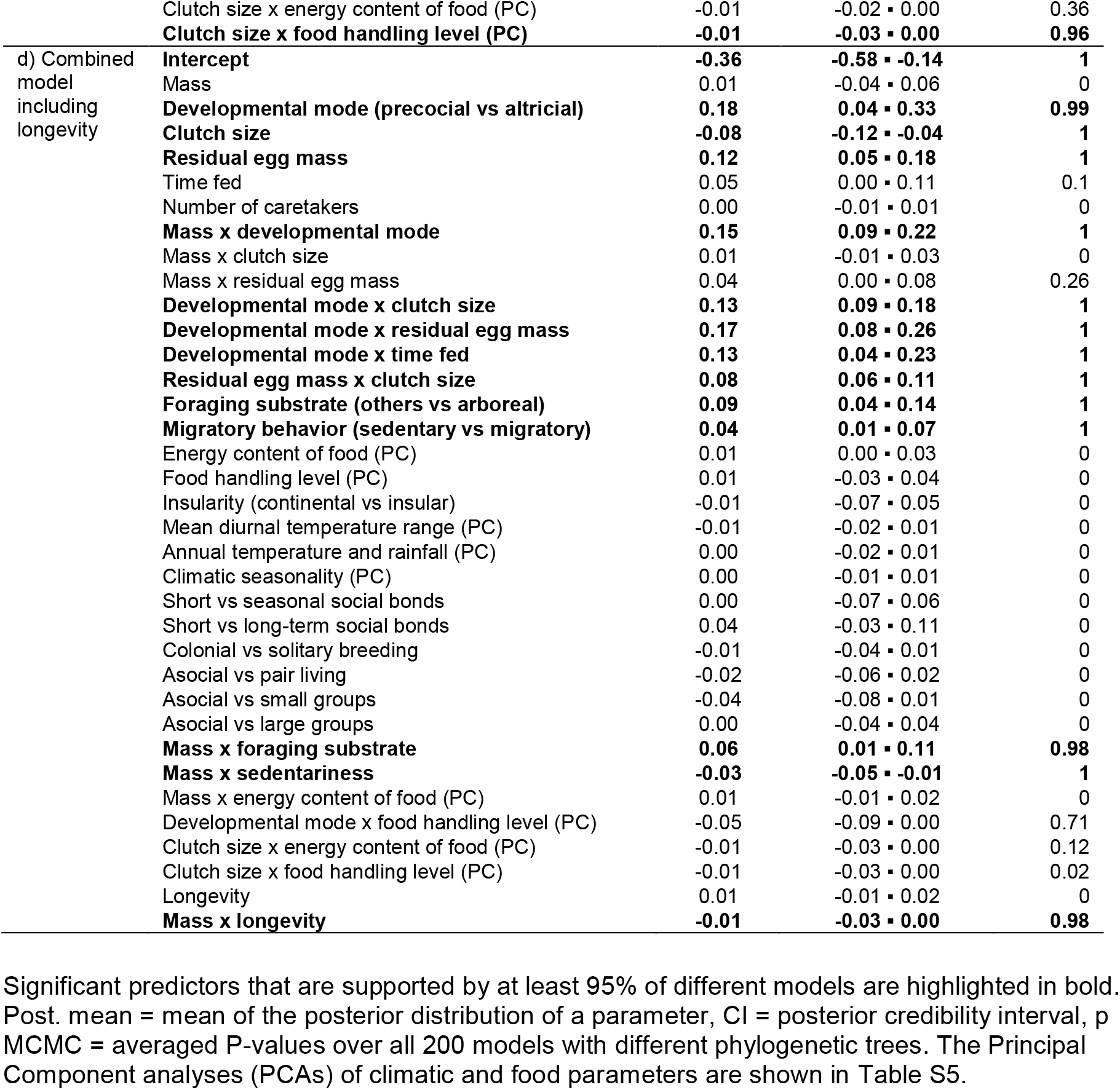
Phylogenetically controlled mixed models in MCMCglmm (1) assessing the effect on relative brain size in birds (model a-c: N=1,176 species, model d: N= 886 species) of a) parental provisioning predictors, b) eco-social predictors, c) both combined, and d) both combined and including longevity.

**Table S3a-c.**
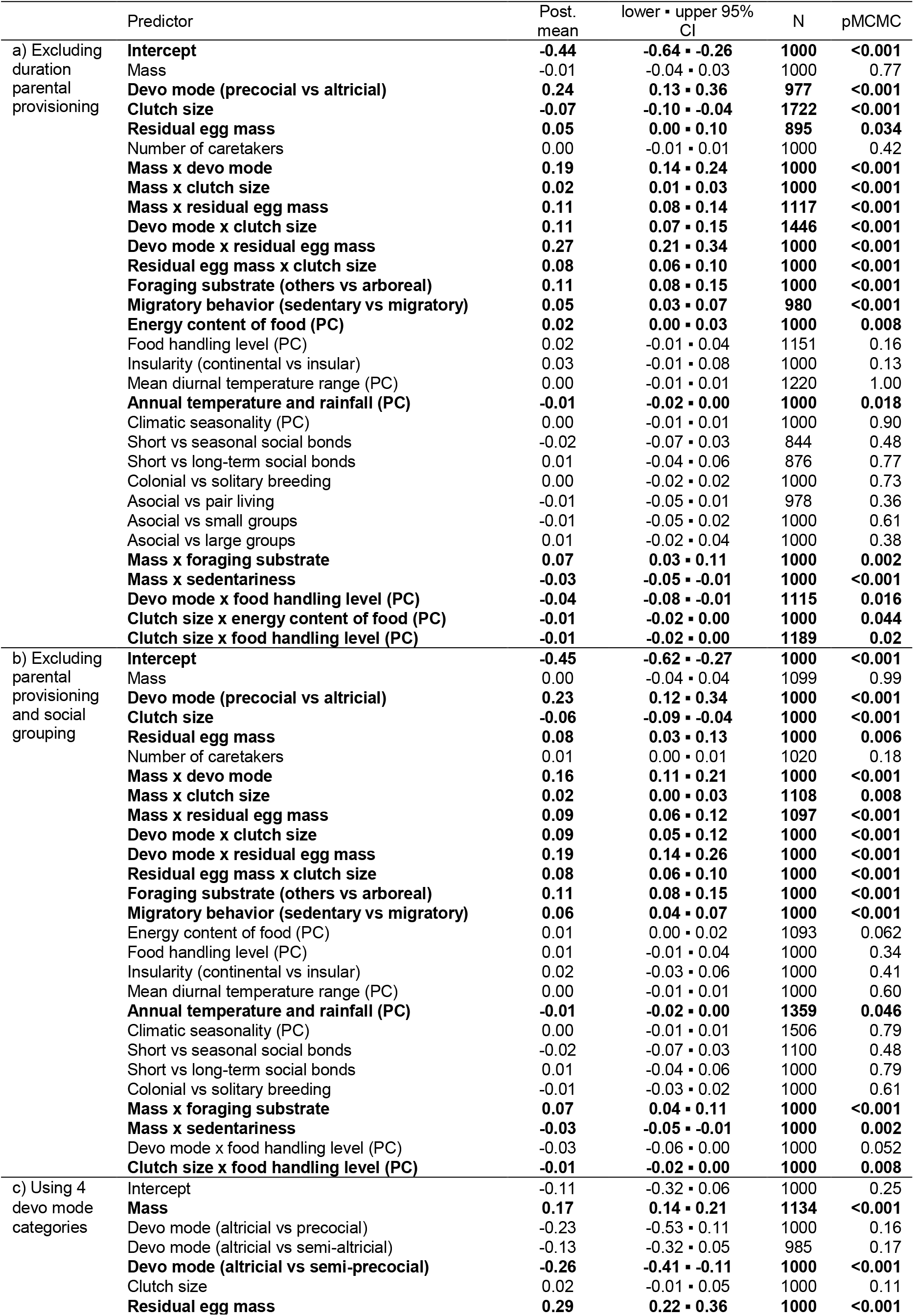
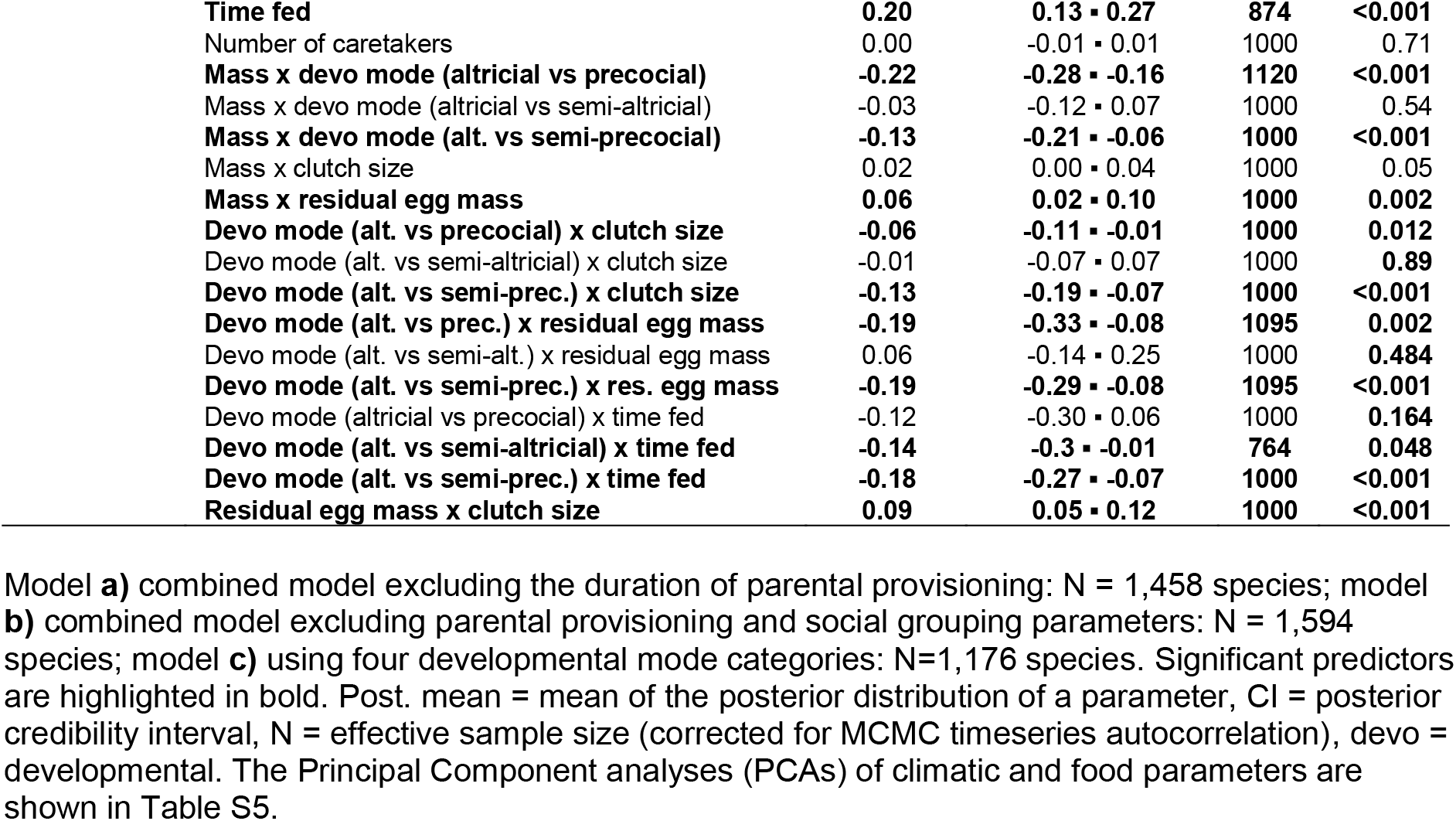
Additional phylogenetically controlled mixed models in MCMCglmm (1) assessing the effect of parental provisioning and eco-social parameters on the evolution of relative brain size in birds.

|  | Predictor | Post.<br>mean | lower • upper 95%<br>CI | N | pMCMC |
| --- | --- | --- | --- | --- | --- |
| a) Excluding duration parental provisioning | **Intercept** | **-0.44** | **-0.64 • -0.26** | **1000** | **<0.001** |
|  | Mass | -0.01 | -0.04 • 0.03 | 1000 | 0.77 |
|  | **Devo mode (precocial vs altricial)** | **0.24** | **0.13 • 0.36** | **977** | **<0.001** |
|  | **Clutch size** | **-0.07** | **-0.10 • -0.04** | **1722** | **<0.001** |
|  | **Residual egg mass** | **0.05** | **0.00 • 0.10** | **895** | **0.034** |
|  | Number of caretakers | 0.00 | -0.01 • 0.01 | 1000 | 0.42 |
|  | **Mass x devo mode** | **0.19** | **0.14 • 0.24** | **1000** | **<0.001** |
|  | **Mass x clutch size** | **0.02** | **0.01 • 0.03** | **1000** | **<0.001** |
|  | **Mass x residual egg mass** | **0.11** | **0.08 • 0.14** | **1117** | **<0.001** |
|  | **Devo mode x clutch size** | **0.11** | **0.07 • 0.15** | **1446** | **<0.001** |
|  | **Devo mode x residual egg mass** | **0.27** | **0.21 • 0.34** | **1000** | **<0.001** |
|  | **Residual egg mass x clutch size** | **0.08** | **0.06 • 0.10** | **1000** | **<0.001** |
|  | **Foraging substrate (others vs arboreal)** | **0.11** | **0.08 • 0.15** | **1000** | **<0.001** |
|  | **Migratory behavior (sedentary vs migratory)** | **0.05** | **0.03 • 0.07** | **980** | **<0.001** |
|  | **Energy content of food (PC)** | **0.02** | **0.00 • 0.03** | **1000** | **0.008** |
|  | Food handling level (PC) | 0.02 | -0.01 • 0.04 | 1151 | 0.16 |
|  | Insularity (continental vs insular) | 0.03 | -0.01 • 0.08 | 1000 | 0.13 |
|  | Mean diurnal temperature range (PC) | 0.00 | -0.01 • 0.01 | 1220 | 1.00 |
|  | **Annual temperature and rainfall (PC)** | **-0.01** | **-0.02 • 0.00** | **1000** | **0.018** |
|  | Climatic seasonality (PC) | 0.00 | -0.01 • 0.01 | 1000 | 0.90 |
|  | Short vs seasonal social bonds | -0.02 | -0.07 • 0.03 | 844 | 0.48 |
|  | Short vs long-term social bonds | 0.01 | -0.04 • 0.06 | 876 | 0.77 |
|  | Colonial vs solitary breeding | 0.00 | -0.02 • 0.02 | 1000 | 0.73 |
|  | Asocial vs pair living | -0.01 | -0.05 • 0.01 | 978 | 0.36 |
|  | Asocial vs small groups | -0.01 | -0.05 • 0.02 | 1000 | 0.61 |
|  | Asocial vs large groups | 0.01 | -0.02 • 0.04 | 1000 | 0.38 |
|  | **Mass x foraging substrate** | **0.07** | **0.03 • 0.11** | **1000** | **0.002** |
|  | **Mass x sedentariness** | **-0.03** | **-0.05 • -0.01** | **1000** | **<0.001** |
|  | **Devo mode x food handling level (PC)** | **-0.04** | **-0.08 • -0.01** | **1115** | **0.016** |
|  | **Clutch size x energy content of food (PC)** | **-0.01** | **-0.02 • 0.00** | **1000** | **0.044** |
|  | **Clutch size x food handling level (PC)** | **-0.01** | **-0.02 • 0.00** | **1189** | **0.02** |
| b) Excluding parental provisioning and social grouping | **Intercept** | **-0.45** | **-0.62 • -0.27** | **1000** | **<0.001** |
|  | Mass | 0.00 | -0.04 • 0.04 | 1099 | 0.99 |
|  | **Devo mode (precocial vs altricial)** | **0.23** | **0.12 • 0.34** | **1000** | **<0.001** |
|  | **Clutch size** | **-0.06** | **-0.09 • -0.04** | **1000** | **<0.001** |
|  | **Residual egg mass** | **0.08** | **0.03 • 0.13** | **1000** | **0.006** |
|  | Number of caretakers | 0.01 | 0.00 • 0.01 | 1020 | 0.18 |
|  | **Mass x devo mode** | **0.16** | **0.11 • 0.21** | **1000** | **<0.001** |
|  | **Mass x clutch size** | **0.02** | **0.00 • 0.03** | **1108** | **0.008** |
|  | **Mass x residual egg mass** | **0.09** | **0.06 • 0.12** | **1097** | **<0.001** |
|  | **Devo mode x clutch size** | **0.09** | **0.05 • 0.12** | **1000** | **<0.001** |
|  | **Devo mode x residual egg mass** | **0.19** | **0.14 • 0.26** | **1000** | **<0.001** |
|  | **Residual egg mass x clutch size** | **0.08** | **0.06 • 0.10** | **1000** | **<0.001** |
|  | **Foraging substrate (others vs arboreal)** | **0.11** | **0.08 • 0.15** | **1000** | **<0.001** |
|  | **Migratory behavior (sedentary vs migratory)** | **0.06** | **0.04 • 0.07** | **1000** | **<0.001** |
|  | Energy content of food (PC) | 0.01 | 0.00 • 0.02 | 1093 | 0.062 |
|  | Food handling level (PC) | 0.01 | -0.01 • 0.04 | 1000 | 0.34 |
|  | Insularity (continental vs insular) | 0.02 | -0.03 • 0.06 | 1000 | 0.41 |
|  | Mean diurnal temperature range (PC) | 0.00 | -0.01 • 0.01 | 1000 | 0.60 |
|  | **Annual temperature and rainfall (PC)** | **-0.01** | **-0.02 • 0.00** | **1359** | **0.046** |
|  | Climatic seasonality (PC) | 0.00 | -0.01 • 0.01 | 1506 | 0.79 |
|  | Short vs seasonal social bonds | -0.02 | -0.07 • 0.03 | 1100 | 0.48 |
|  | Short vs long-term social bonds | 0.01 | -0.04 • 0.06 | 1000 | 0.79 |
|  | Colonial vs solitary breeding | -0.01 | -0.03 • 0.02 | 1000 | 0.61 |
|  | **Mass x foraging substrate** | **0.07** | **0.04 • 0.11** | **1000** | **<0.001** |
|  | **Mass x sedentariness** | **-0.03** | **-0.05 • -0.01** | **1000** | **0.002** |
|  | **Devo mode x food handling level (PC)** | **-0.03** | **-0.06 • 0.00** | **1000** | **0.052** |
|  | **Clutch size x food handling level (PC)** | **-0.01** | **-0.02 • 0.00** | **1000** | **0.008** |
| c) Using 4 devo mode categories | **Intercept** | **-0.11** | **-0.32 • 0.06** | **1000** | **0.25** |
|  | **Mass** | **0.17** | **0.14 • 0.21** | **1134** | **<0.001** |
|  | Devo mode (altricial vs precocial) | -0.23 | -0.53 • 0.11 | 1000 | 0.16 |
|  | Devo mode (altricial vs semi-altricial) | -0.13 | -0.32 • 0.05 | 985 | 0.17 |
|  | **Devo mode (altricial vs semi-precocial)** | **-0.26** | **-0.41 • -0.11** | **1000** | **<0.001** |
|  | Clutch size | 0.02 | -0.01 • 0.05 | 1000 | 0.11 |
|  | **Residual egg mass** | **0.29** | **0.22 • 0.36** | **1000** | **<0.001** |

|  |  |  |  |  |
| --- | --- | --- | --- | --- |
| **Time fed** | **0.20** | **0.13 ▪ 0.27** | **874** | **<0.001** |
| Number of caretakers | 0.00 | -0.01 ▪ 0.01 | 1000 | 0.71 |
| **Mass x devo mode (altricial vs precocial)** | **-0.22** | **-0.28 ▪ -0.16** | **1120** | **<0.001** |
| Mass x devo mode (altricial vs semi-altricial) | -0.03 | -0.12 ▪ 0.07 | 1000 | 0.54 |
| **Mass x devo mode (alt. vs semi-precocial)** | **-0.13** | **-0.21 ▪ -0.06** | **1000** | **<0.001** |
| Mass x clutch size | 0.02 | 0.00 ▪ 0.04 | 1000 | 0.05 |
| **Mass x residual egg mass** | **0.06** | **0.02 ▪ 0.10** | **1000** | **0.002** |
| **Devo mode (alt. vs precocial) x clutch size** | **-0.06** | **-0.11 ▪ -0.01** | **1000** | **0.012** |
| Devo mode (alt. vs semi-altricial) x clutch size | -0.01 | -0.07 ▪ 0.07 | 1000 | **0.89** |
| **Devo mode (alt. vs semi-prec.) x clutch size** | **-0.13** | **-0.19 ▪ -0.07** | **1000** | **<0.001** |
| **Devo mode (alt. vs prec.) x residual egg mass** | **-0.19** | **-0.33 ▪ -0.08** | **1095** | **0.002** |
| Devo mode (alt. vs semi-alt.) x residual egg mass | 0.06 | -0.14 ▪ 0.25 | 1000 | **0.484** |
| **Devo mode (alt. vs semi-prec.) x res. egg mass** | **-0.19** | **-0.29 ▪ -0.08** | **1095** | **<0.001** |
| Devo mode (altricial vs precocial) x time fed | -0.12 | -0.30 ▪ 0.06 | 1000 | **0.164** |
| **Devo mode (alt. vs semi-altricial) x time fed** | **-0.14** | **-0.3 ▪ -0.01** | **764** | **0.048** |
| **Devo mode (alt. vs semi-prec.) x time fed** | **-0.18** | **-0.27 ▪ -0.07** | **1000** | **<0.001** |
| **Residual egg mass x clutch size** | **0.09** | **0.05 ▪ 0.12** | **1000** | **<0.001** |

**Table S4.** Additional phylogenetically controlled mixed model in MCMCglmm (1) assessing the effect of parental provisioning and eco-social parameters on the evolution of relative brain size in birds, using lineage specific slopes, following the clade taxonomy by Prum et al. (2).

| Predictor | Post.<br>mean | lower • upper 95%<br>CI | N | pMCMC |
| --- | --- | --- | --- | --- |
| Intercept | 0.00 | -0.19 • 0.15 | 1345 | 0.99 |
| **Mass** | **-0.07** | **-0.10 • -0.03** | **1000** | **<0.001** |
| Developmental mode (precocial vs altricial) | 0.01 | -0.10 • 0.13 | 1000 | 0.86 |
| **Clutch size** | **-0.06** | **-0.08 • -0.02** | **745** | **<0.001** |
| **Residual egg mass** | **0.13** | **0.08 • 0.18** | **1000** | **<0.001** |
| Time fed | 0.01 | -0.03 • 0.06 | 861 | 0.73 |
| Number of caretakers | 0.00 | -0.01 • 0.01 | 1000 | 0.78 |
| **Mass x developmental mode** | **0.09** | **0.04 • 0.14** | **1000** | **<0.001** |
| Mass x clutch size | 0.01 | -0.01 • 0.02 | 1000 | 0.24 |
| Mass x residual egg mass | 0.01 | -0.03 • 0.04 | 1000 | 0.70 |
| **Developmental mode x clutch size** | **0.09** | **0.05 • 0.12** | **1141** | **<0.001** |
| **Developmental mode x residual egg mass** | **0.12** | **0.04 • 0.19** | **1000** | **0.004** |
| **Developmental mode x time fed** | **0.13** | **0.05 • 0.20** | **1000** | **<0.001** |
| **Residual egg mass x clutch size** | **0.07** | **0.05 • 0.09** | **1000** | **<0.001** |
| **Foraging substrate (others vs arboreal)** | **0.05** | **0.01 • 0.10** | **1000** | **0.008** |
| **Migratory behavior (sedentary vs migratory)** | **0.05** | **0.03 • 0.07** | **1148** | **<0.001** |
| Energy content of food (PC) | 0.01 | 0.00 • 0.02 | 1000 | 0.33 |
| Food handling level (PC) | 0.01 | -0.02 • 0.04 | 1000 | 0.61 |
| Insularity (continental vs insular) | 0.00 | -0.05 • 0.04 | 1000 | 0.87 |
| Mean diurnal temperature range (PC) | 0.00 | -0.01 • 0.01 | 1000 | 0.45 |
| Annual temperature and rainfall (PC) | 0.00 | -0.02 • 0.01 | 757 | 0.48 |
| Climatic seasonality (PC) | 0.00 | 0.00 • 0.01 | 839 | 0.39 |
| Short vs seasonal social bonds | -0.01 | -0.06 • 0.04 | 1000 | 0.75 |
| Short vs long-term social bonds | 0.02 | -0.02 • 0.08 | 1000 | 0.36 |
| Colonial vs solitary breeding | -0.01 | -0.04 • 0.01 | 1000 | 0.32 |
| Asocial vs pair living | -0.02 | -0.05 • 0.01 | 1000 | 0.26 |
| Asocial vs small groups | -0.02 | -0.05 • 0.02 | 1000 | 0.43 |
| Asocial vs large groups | 0.00 | -0.03 • 0.03 | 1054 | 0.86 |
| Mass x foraging substrate | 0.00 | -0.04 • 0.04 | 892 | 0.90 |
| **Mass x sedentariness** | **-0.03** | **-0.05 • -0.01** | **1000** | **0.004** |
| Mass x energy content of food (PC) | 0.01 | -0.01 • 0.02 | 1000 | 0.37 |
| **Developmental mode x food handling level (PC)** | **-0.04** | **-0.07 • 0.00** | **1000** | **0.038** |
| Clutch size x energy content of food (PC) | -0.01 | -0.02 • 0.01 | 1000 | 0.39 |
| Clutch size x food handling level (PC) | -0.01 | -0.02 • 0.00 | 1000 | 0.12 |

**Table S5.** Principal Component Analyses (PCA) loadings and variance explained. Loadings of the main contributors to the different component are highlighted in bold.

| **a) Food PCA** | PC Food1 | PC Food2 |
| --- | --- | --- |
| Food energy content / 100g | 0.12 | **0.99** |
| Food fiber content / 100g | **-0.96** | -0.04 |
| Food handling levels | **0.95** | 0.17 |
| SS loadings | 1.84 | 1.01 |
| Proportion Variance | 0.61 | 0.34 |
| Cumulative Variance | 0.61 | 0.95 |
| Proportion Explained | 0.64 | 0.36 |
| Cumulative Proportion | 0.64 | 1.00 |

| **b) Climatic PCA** | PC Climate1 | PC Climate2 | PC Climate3 |
| --- | --- | --- | --- |
| Bio1_mean Annual Mean Temperature | **0.89** | 0.06 | 0.04 |
| Bio1_stdev | **-0.62** | **0.53** | 0.04 |
| Bio2_mean Mean Diurnal Range | -0.14 | 0.19 | **0.81** |
| Bio2_stdev | -0.30 | **0.71** | 0.43 |
| Bio3_mean Isothermality | **0.89** | -0.02 | -0.22 |
| Bio3_stdev | -0.02 | 0.87 | 0.10 |
| Bio4_mean Temperature Seasonality | **-0.96** | 0.02 | 0.13 |
| Bio4_stdev | **-0.78** | 0.50 | 0.14 |
| Bio7_mean Temperature Annual Range | **-0.90** | 0.08 | 0.34 |
| Bio7_stdev | **-0.69** | **0.61** | 0.22 |
| Bio12_mean Annual Precipitation | **0.56** | -0.01 | **-0.73** |
| Bio12_stdev | 0.28 | 0.65 | -0.44 |
| Bio15_mean Precipitation Seasonality | 0.01 | 0.17 | **0.80** |
| Bio15_stdev | -0.11 | 0.74 | 0.39 |
| SS loadings | 5.32 | 3.22 | 2.63 |
| Proportion Variance | 0.38 | 0.23 | 0.19 |
| Cumulative Variance | 0.38 | 0.61 | 0.80 |
| Proportion Explained | 0.48 | 0.29 | 0.24 |
| Cumulative Proportion | 0.48 | 0.76 | 1.00 |

**a)** Food items PCA. **b)** Climatic PCA.

**S6 Table.** Variance inflation factor (VIF) analyses of all parameters included in the combined model.

| Predictor | VIF |
| --- | --- |
| Mass | 1.188 |
| Developmental mode (precocial vs altricial) | 1.162 |
| Clutch size | 1.219 |
| Residual egg mass | 1.273 |
| Time fed | 1.906 |
| Number of caretakers | 1.167 |
| Foraging substrate (others vs arboreal) | 1.075 |
| Migratory behavior (sedentary vs migratory) | 1.352 |
| Energy content of food (PC) | 1.095 |
| Food handling level (PC) | 1.135 |
| Insularity (continental vs insular) | 1.564 |
| Mean diurnal temperature range (PC) | 1.600 |
| Annual temperature and rainfall (PC) | 2.076 |
| Climatic seasonality (PC) | 1.108 |
| Social bonds | 1.567 |
| Colonial vs solitary breeding | 1.031 |
| Grouping pattern | 1.073 |

Variance inflation factors (VIF) were calculated from a Bayesian phylogenetically controlled mixed model including parental provisioning and eco-social predictors. VIF is assumed to be the term inflating the regression coefficient's variance estimate $\widehat{var}(\beta_i) = \left( \frac{s^2}{(n-1)\widehat{var}(X_i)} \right) \frac{1}{1-R_i^2} = \left( \frac{s^2}{(n-1)\widehat{var}(X_i)} \right) VIF_i$ , where $s^2$ is the root mean squared error of regression, $n$ is the sample size, $X_i$ is the $i$ -th column of the design matrix. $R_i^2$ then becomes the GLMM $R^2$ analogue for the model: $X_i = \alpha_0 + \alpha_1 X_1 + \dots + \alpha_{i-1} X_{i-1} + \alpha_{i+1} X_{i+1} + \dots + \alpha_k X_k$ .

**S7 Table.**
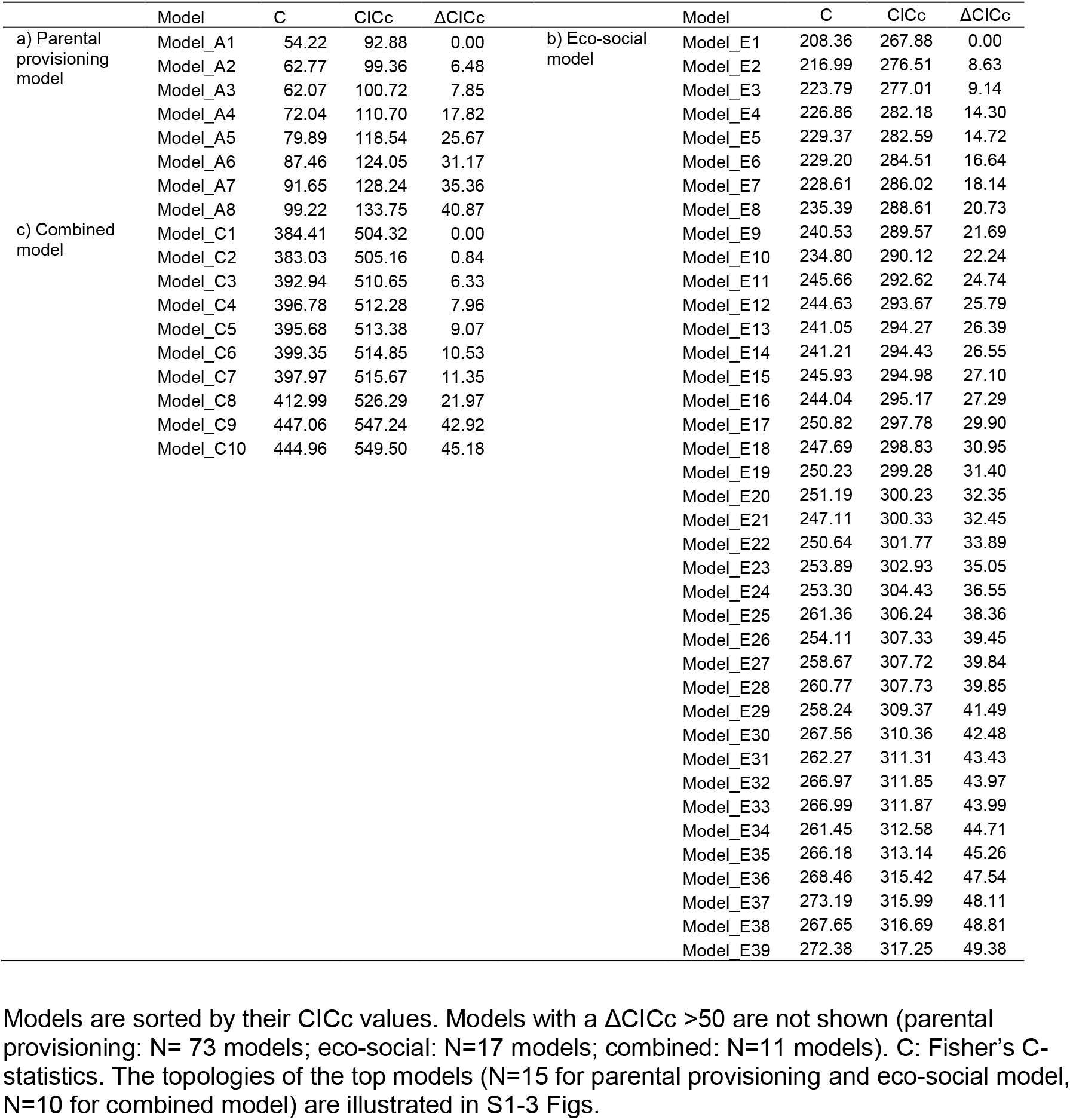
Model comparisons of the models included in the phylogenetic d-separation path analyses.

**Dataset S1 (separate file)**. Dataset used for analyses.

## Notes

### Competing Interest Statement

The authors have declared no competing interest.

### Summary of Updates

We explain now that larger brains only can evolve if there is both a benefit and individuals can afford the energetic costs of growing and maintaining them. We also clarify a number of methodological issues, and improve our wording with respect to causality.

